# Akt1-associated actomyosin remodelling is required for nuclear lamina dispersal and nuclear shrinkage in epidermal terminal differentiation

**DOI:** 10.1101/868034

**Authors:** Clare Rogerson, Duncan Wotherspoon, Ryan F L O’Shaughnessy

## Abstract

Keratinocyte cornification and epidermal barrier formation are tightly controlled processes, which require complete degradation of intracellular organelles, including removal of keratinocyte nuclei. Keratinocyte nuclear destruction requires Akt1-dependent phosphorylation and degradation of the nuclear lamina protein, Lamin A/C, essential for nuclear integrity. However, the molecular mechanisms that result in complete nuclear removal and their regulation are not well defined. Post-confluent cultures of rat epidermal keratinocytes (REKs) undergo spontaneous and complete differentiation, allowing visualisation and perturbation of the differentiation process *in vitro*. We demonstrate that there is dispersal of phosphorylated Lamin A/C to structures throughout the cytoplasm in differentiating keratinocytes. We show that the dispersal of phosphorylated Lamin A/C is Akt1-dependent and these structures are specific for the removal of Lamin A/C from the nuclear lamina; nuclear contents and Lamin B were not present in these structures. Immunoprecipitation identified a group of functionally related Akt1 target proteins involved in Lamin A/C dispersal, including actin, which forms cytoskeletal microfilaments, Arp3, required for actin filament nucleation, and Myh9, a component of myosin IIa, a molecular motor that can translocate along actin filaments. Disruption of actin filament polymerisation, nucleation or myosin IIa activity prevented formation and dispersal of cytoplasmic Lamin A/C structures. Live imaging of keratinocytes expressing fluorescently tagged nuclear proteins showed a nuclear volume reduction step taking less than 40 minutes precedes final nuclear destruction. Preventing Akt1-dependent Lamin A/C phosphorylation and disrupting cytoskeletal Akt1-associated proteins prevented nuclear volume reduction. Single cell RNA sequencing of differentiating keratinocytes identified gene changes correlated with lamin dispersal, which we propose are due to changes in lamina-associated domains upon Lamin A/C dispersal. We propose keratinocyte nuclear destruction and differentiation requires myosin II activity and the actin cytoskeleton for two intermediate processes: Lamin A/C dispersal and rapid nuclear volume reduction.

## Introduction

The mammalian epidermis is an essential barrier between an organism and its environment (Madison 2003; Eckhart and Zeeuwen 2018). The epidermis has four layers of keratinocytes: basal, spinous, granular and cornified layers, the last of which is comprised of corneocytes (Watt 1989; Eckhart et al. 2013). Formation of a healthy skin barrier requires continual differentiation of epidermal keratinocytes into corneocytes, (Eckhart et al. 2013). The transition of keratinocytes from granular layer cells to corneocytes is a highly controlled and irreversible terminal process. As part of this, keratinocytes undergo major morphological changes, involving removal of all their intracellular organelles, including their nuclei, allowing corneocytes to contain an extremely high proportion of keratin and form the rigid outer epidermal cell layer (Eckhart et al. 2013; Matsui and Amagai 2015).

The disruption of nuclear degradation leading to retention of nuclei in the cornified layer, known as parakeratosis, is a frequent observation in many skin diseases (Song and Shea 2010; Cardoso et al. 2017); however, the mechanisms by which keratinocytes remove their nuclei are not well defined (Rogerson, Bergamaschi, and O’Shaughnessy 2018; Eckhart et al. 2013). Nuclear removal is dependent on the Akt1 kinase. Akt1 shRNA knockdown decreased phosphorylation of the nuclear lamina component Lamin A/C, decreased Lamin A/C degradation, and increased nuclear retention in the cornified layer (O’Shaughnessy et al. 2007; Naeem et al. 2015). Subsequent to nuclear lamina breakdown, DNA degrading enzymes are also required for nuclear content breakdown; murine epidermis lacking DNase1L2 and DNase2 is parakeratotic, however the parakeratotic nuclei show removal of Lamin A/C indicating DNA degradation occurs after lamina removal (Fischer et al. 2017). Proteins important in autophagic processes are also required for removal of nuclear content; dysregulation of autophagy-related proteins correlates with decreased nuclear degradation and the autophagic marker LC3 localises close to the nucleus in differentiating cells (Akinduro et al. 2016; Park et al. 2009).

The transition from granular keratinocytes to corneocytes occurs over a 24 hour period, with the nuclear removal process estimated to take 6 hours (Eckhart et al. 2013; Lavker and Matoltsy 1970). However, the mechanisms preceding removal of the nucleus are currently unidentified (Rogerson, Bergamaschi, and O’Shaughnessy 2018). Using rat epidermal keratinocytes (REKs), which spontaneously differentiate in submerged culture, we have followed nuclear degradation in an *in vitro* model without forced initiation of differentiation by calcium switch. We identified Akt1-dependent dispersion of nuclear lamina components throughout the cytoplasm in differentiating keratinocytes that precedes nuclear degradation. Following fluorescently tagged nuclear proteins in REK differentiation in real time has allowed us to identify, for the first time, rapid nuclear shrinkage as an important nuclear degradation intermediate. We also isolated a ‘degradosome’ complex of proteins that associate with Akt1, including actin and the actin-binding proteins Myh9 and Arp3 that are involved in nuclear removal. Treatments that affect lamina dispersal also identified gene expression changes that may be controlled by lamina-associated domain removal. These results provide evidence for two nuclear degradation intermediates in keratinocyte differentiation and the characterisation of a protein complex important for nuclear degradation regulation.

## Results

### Akt1-dependent dispersal of phosphorylated lamina proteins in keratinocyte differentiation

Akt1 is required for Lamin A/C phosphorylation and Akt1 knockdown prevents Lamin A/C degradation and nuclear removal, suggesting Lamin A/C phosphorylation is important for nuclear lamina breakdown in keratinocyte nuclear removal (Naeem et al. 2015). Phosphorylated serine 404 (pSer404) Lamin A/C disperses throughout granular layer cells in human and murine skin sections and REK organotypic cultures (Naeem et al. 2015). We determined that pSer404 Lamin A/C dispersal occurred in suprabasal, differentiating REKs in monolayer culture, (Figure 1A and B). Akt1 co-localised with pSer404 Lamin A/C at the nucleus and at dispersed structures throughout the cytoplasm, (Figure 1C). Lamin A/C dispersal was reduced in Akt1 knockdown REK organotypic cultures (Naeem et al. 2015), and suprabasal Akt1 knockdown REKs had increased nuclear size in post-confluent cultures compared to controls, (Naeem et al. 2015), (Figure 1D), and these larger nuclei-containing cells are larger and less granular based on flow cytometry side scatter (SSC) measurements, (Figure 1E-H). Conversely, rapamycin treatment, which increases Akt1 activity (Sully et al. 2013) increased the number of cells with Lamin A/C dispersal, (Figure 1I and J). Akt1 activity is therefore required for Lamin A/C dispersal and nuclear degradation.

**Figure 1.**
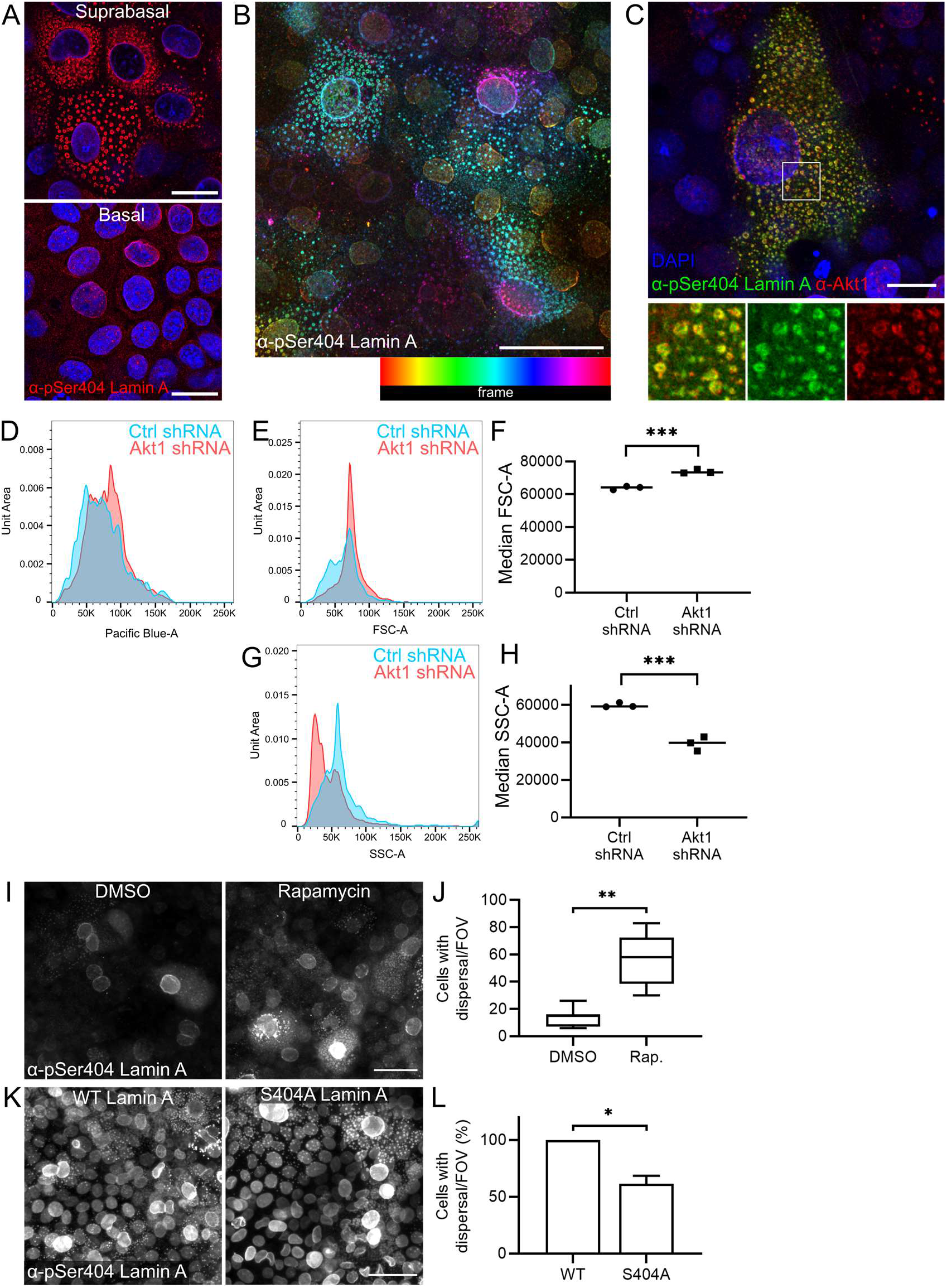
Phosphorylated Lamin A in keratinocyte differentiation and nuclear degradation. A – pSer404 Lamin A/C staining in single confocal sections of basal and suprabasal REKs. Scale bars = 20 μm. B - pSer404 Lamin A/C signal colour-coded according to the frame of the z-stack. Yellow = basal signal, blue and pink = more suprabasal signal. Scale bar = 20 μm. C - Akt1 co-localises with pSer404 Lamin A/C in a single confocal section of suprabasal REKs. Box indicates enlarged area, scale bar = 20 μm. D – Area of Hoechst 33342 staining (nuclear content) of control or Akt1 knockdown large differentiating REKs. E – FSC-A (Area) area of control or Akt1 knockdown large differentiating REKs. F – Median FSC-A in control or Akt1 knockdown large differentiating REKs, in triplicate *** p ≤ 0.001. G ‒SSC-A (Area) of control or Akt1 knockdown large differentiating REKs. H – Median SSC-A in control or Akt1 knockdown large differentiating REKs, in triplicate *** p ≤ 0.001. I – DMSO or Rapamycin treated post-confluent REKs stained for pSer404 Lamin A/C. Maximum projections of confocal z-stacks, scale bar = 50 μm. J - Number of cells with dispersed pSer404 Lamin A/C per field of view (FOV), ≥5 FOV, Welch’s t test, ** p ≤ 0.01. K – REKs expressing WT or S404A Lamin A stained for pSer404 Lamin A/C. Epifluorescence image, scale bar = 50 μm. L - Number of cells with dispersed pSer404 Lamin A/C per FOV compared to WT Lamin A. Two independent experiments, three FOV per experiment, unpaired t-test, * p ≤ 0.05.

Overexpression of ‘non-AKT-phosphorylatable’ S404A Lamin A decreased Lamin A/C dispersal, (Figure 1K and L), and reduced expression of the keratinocyte differentiation marker loricrin, (Supplementary Figure 1A and B). This suggested that Akt1-dependent phosphorylation of Lamin A/C also affected differentiation. Overexpression of ‘phosphomimetic’ S404D Lamin A did not increase Lamin A/C dispersal, nor did it increase Loricrin staining or alter nuclear size, (Supplementary Figure 1A-E). This construct did have a lower transfection efficiency, Supplementary Figure 1F, suggesting an intolerance to the expression of the phosphomimetic construct, or alternatively, the majority of Lamin A/C is phosphorylated, and so the system is not affected by the phosphomimetic.

### Lamina dispersal is specific to Lamin A/C

Dispersed pSer404 Lamin A/C structures did not contain any identifiable nuclear content; they did not consistently stain with the DNA intercalating dye DAPI and did not co-localise with Histone proteins, either endogenous Histone H3 or H2B or an overexpressed Histone H2B-mCherry, (Figure 2A-C, Supplementary Figure 1G). Ran, a nuclear transport protein, dispersed to pSer404 Lamin A/C cytoplasmic structures, (Figure 2D), indicating their origin at the nucleus. However, the structures did not contain mCherry targeted to the nucleus with a nuclear localisation sequence (NLS-mCherry), (Figure 1E); indicating these structures did not contain functional nuclear pores. Additionally, Lamin B1 did not disperse to cytoplasmic Akt1-containing structures, (Figure 2F); suggesting Lamin A/C but not Lamin B1 is involved in this process.

**Figure 2.**
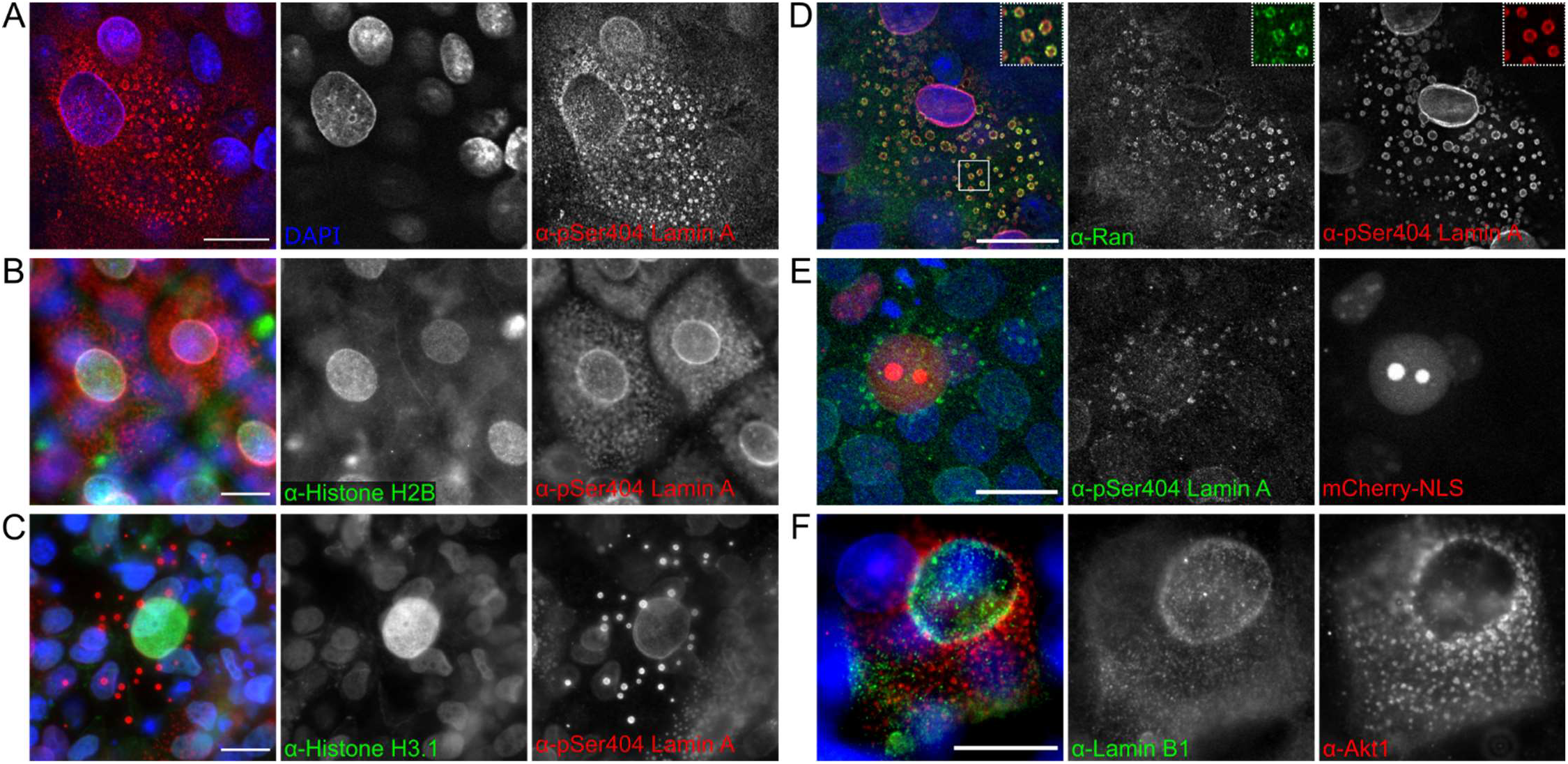
Lamina protein dispersal to the cytoplasm is pSer404 Lamin A/C specific and the cytoplasmic structures do not contain histone proteins, NLS-targeted protein or Lamin B1. A – pSer404 Lamin A/C in suprabasal REKs co-stained with DAPI (DNA marker). Confocal z-section. B-C - pSer404 Lamin A/C in suprabasal REKs co-stained for Histone H2B (B) and Histone H3 (C). Epifluorescence images. D – pSer404 Lamin A/C in suprabasal REKs co-stained for Ran GTPase. Confocal z-section, boxed area is enlarged. E - NLS-mCherry expressing REK co-stained with pSer404 Lamin A/C in suprabasal REKs. Maximum projection of confocal z-sections. F - Lamin B1 in suprabasal REKs co-stained with Akt1. Epifluorescence image Scale bars = 20 μm.

### Akt1-associated proteins form a ‘degradosome’ complex in differentiating REKs

AKT family kinases have a large number of downstream target proteins involved in a wide range of cellular processes that may be tissue, cell and cellular location specific (Manning and Cantley 2007), suggesting the potential of novel Akt1-interacting proteins being involved in lamin dispersal and nuclear shrinkage. To explore this we performed immunoprecipitation with an Akt1-specific antibody, followed by mass spectrometric analysis of eight major SDS-PAGE gel bands. Excluding mitochondrial and keratin proteins, we identified 21 potential Akt1-interacting partners in REKs (Figure 3A and Supplementary Figure 2A), including the previously identified Akt1 target, HspB1 (O’Shaughnessy et al. 2007). STRING analysis of these candidates identified several previously identified interactions or links between these proteins, (Figure 3B). Co-staining with Akt1 or pSer404 Lamin A antibodies demonstrated co-localisation or adjacent expression on cytoplasmic structures for filamentous actin (phalloidin), Arp3, Cops4, Myh9, Ran, Jup and Stambp, (Figures 2D and 3C-H). Staining of mouse epidermis indicated co-localisation of dispersed Akt1 or pSer404 Lamin A staining with Myh9, Jup, Ran and Stambp, (Figure 4A-D). Immunoprecipitation confirmed β-actin, Arp3, and Myh9 interacted with Akt1 (Figure 4E). These results suggest functionally related Akt1 interactors suggestive of a ‘degradosome’ complex including actin and the actin-binding proteins, Myh9 and Arp3, associating with Akt1 and with cytoplasmic pSer404 Lamin A in differentiating REKs, (Figure 4F). Myh9 is a component of myosin IIa, a molecular motor that can translocate along actin filaments, and Arp3 is a component of the Arp2/3 complex required for actin filament nucleation (Marigo et al. 2004; Mullins, Heuser, and Pollard 1998). This suggests a dependence on the actin cytoskeleton and actin-binding proteins for Akt1 function in epidermal terminal differentiation.

**Figure 3.**
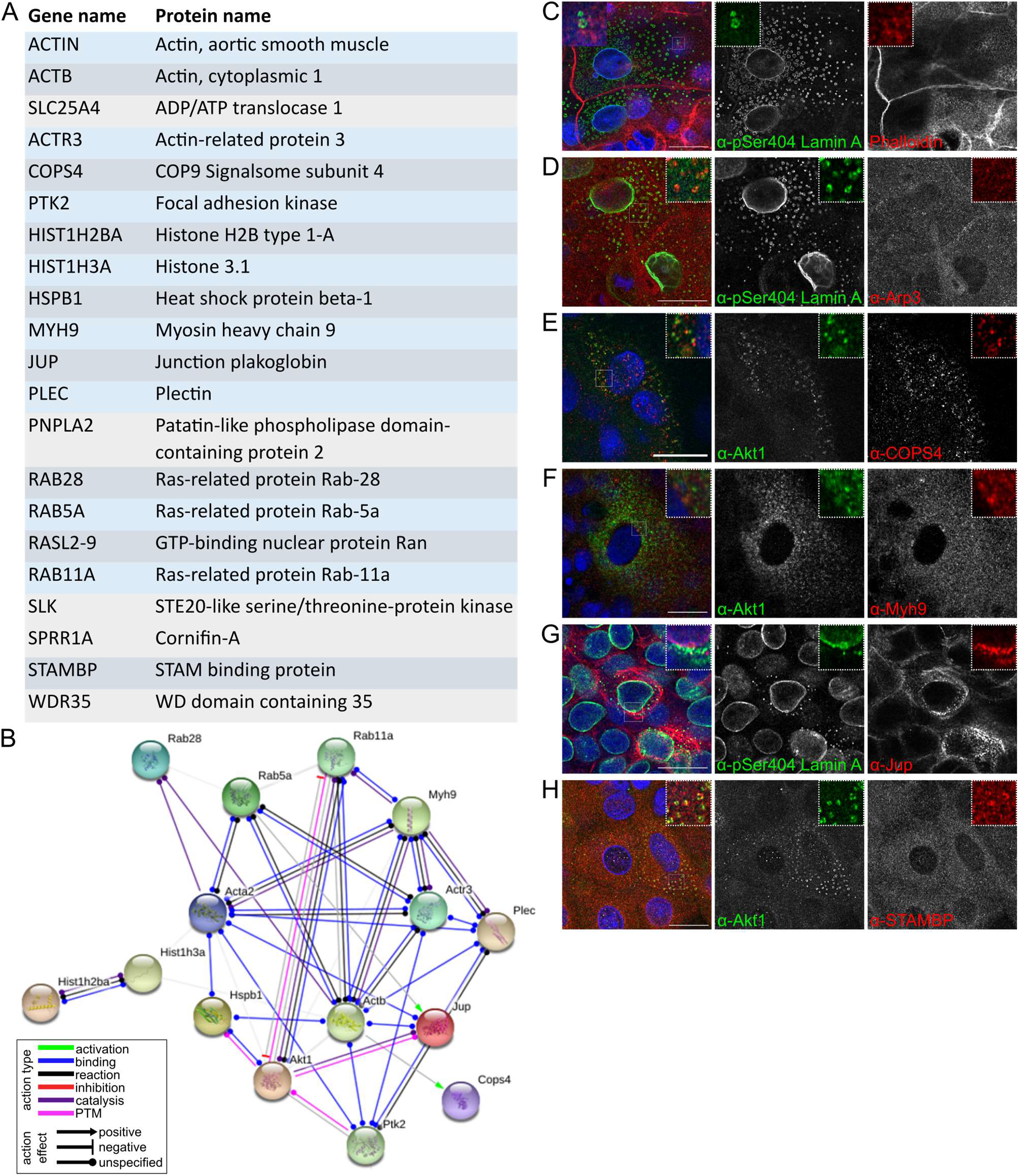
Identification of Akt1 interactors and co-staining with Akt1 and pSer404 Lamin A in differentiating keratinocytes. A – List of Akt1 interactors identified by LC-MS/MS of Akt1 co-immunoprecipitate. B – STRING analysis of interactors in (A). PTM = posttranslational modification. C-H - Suprabasal differentiating REKs with Akt1 or pSer404 Lamin A/C co-stained for actin (phalloidin) (C), Arp3 (D), COPS4 (E), Myh9 (F), Jup (G), STAMBP (H). Confocal z-sections, scale bars = 20 μm.

**Figure 4.**
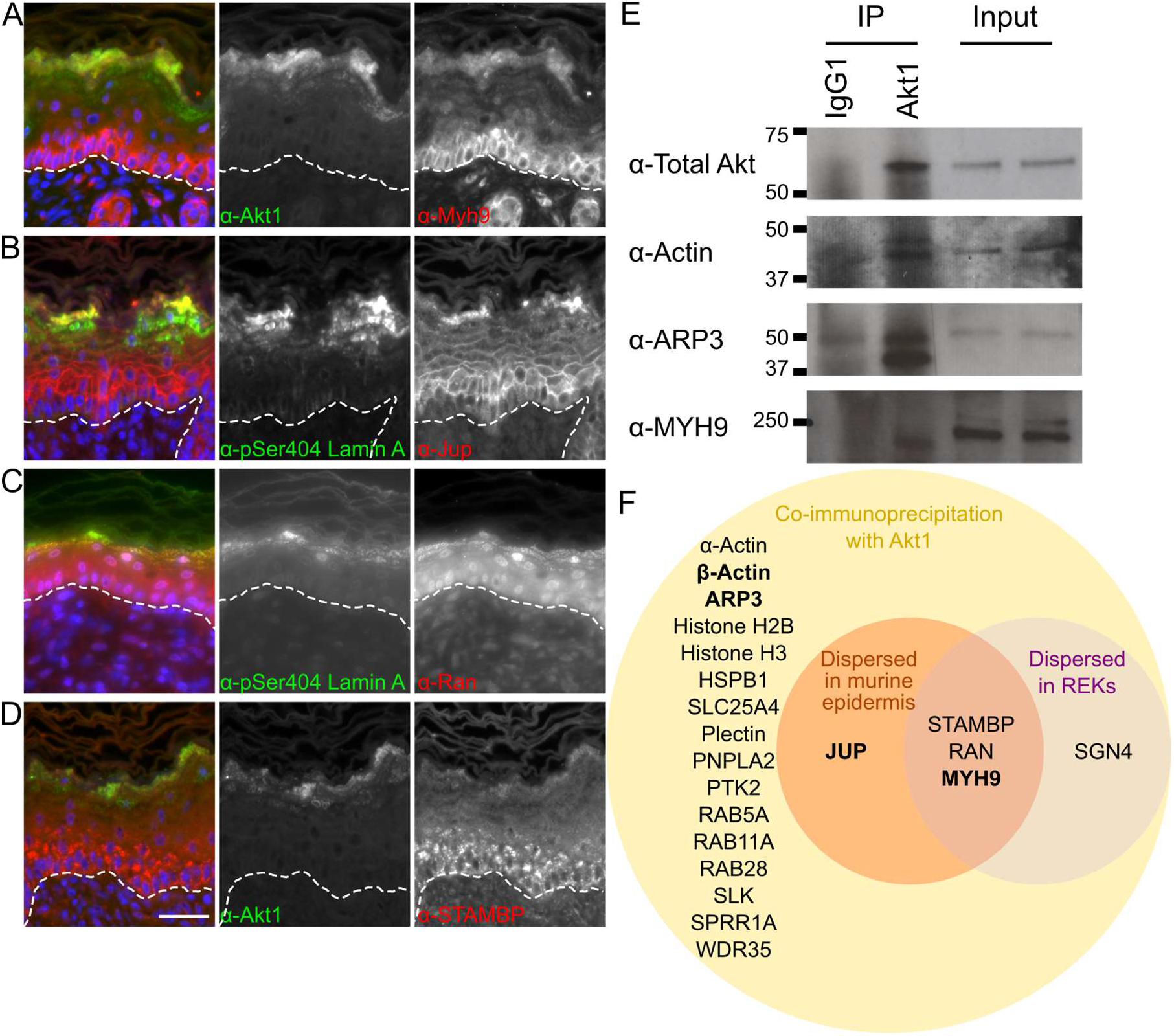
Akt1 interactors in murine epidermis and in Akt1 immunoprecipitate. A-D - Neonatal wild-type murine epidermis stained for Akt1 or pSer404 Lamin A and co-stained for Myh9 (A), Jup (B), Ran (C), STAMBP (D). E – IgG1 and Akt1 immunoprecipitate immunoblotted for actin, Arp3 and Myh9. F – Summary of Akt1 interactors identified by mass spectrometry in an Akt1 co-immunoprecipitate that were also detected to be dispersed with Akt1 and pSer404 Lamin A/C in REKs, murine epidermal sections, and confirmed by western blotting of Akt1 co-immunoprecipitate in bold.

### Akt1-associated actin remodelling is required for nuclear degradation

To test the requirement for actin and myosin II motors in nuclear degradation, we chemically disrupted their function; using latrunculin B to block actin polymerisation and blebbistatin to inhibit myosin II activity in post-confluent REKs. Latrunculin B and blebbistatin treatment inhibited Lamin A/C dispersal to the cytoplasm, (Figure 5A and B), and increased nuclear size in large granular differentiating REKs, (Figure 5D-E and Supplementary Figure 2F-G). These treatments did not affect levels of phosphorylated full length, and cleaved, Lamin A/C in post-confluent REK cultures, (Figure 5C, F and G). Treatment with CK666, an Arp2/3 specific inhibitor decreased phosphorylated Lamin A/C and dispersal at 50 μM, (Figure 5H-J) but had no significant effect on nuclear size, (Supplementary Figure 2H and I).

**Figure 5.**
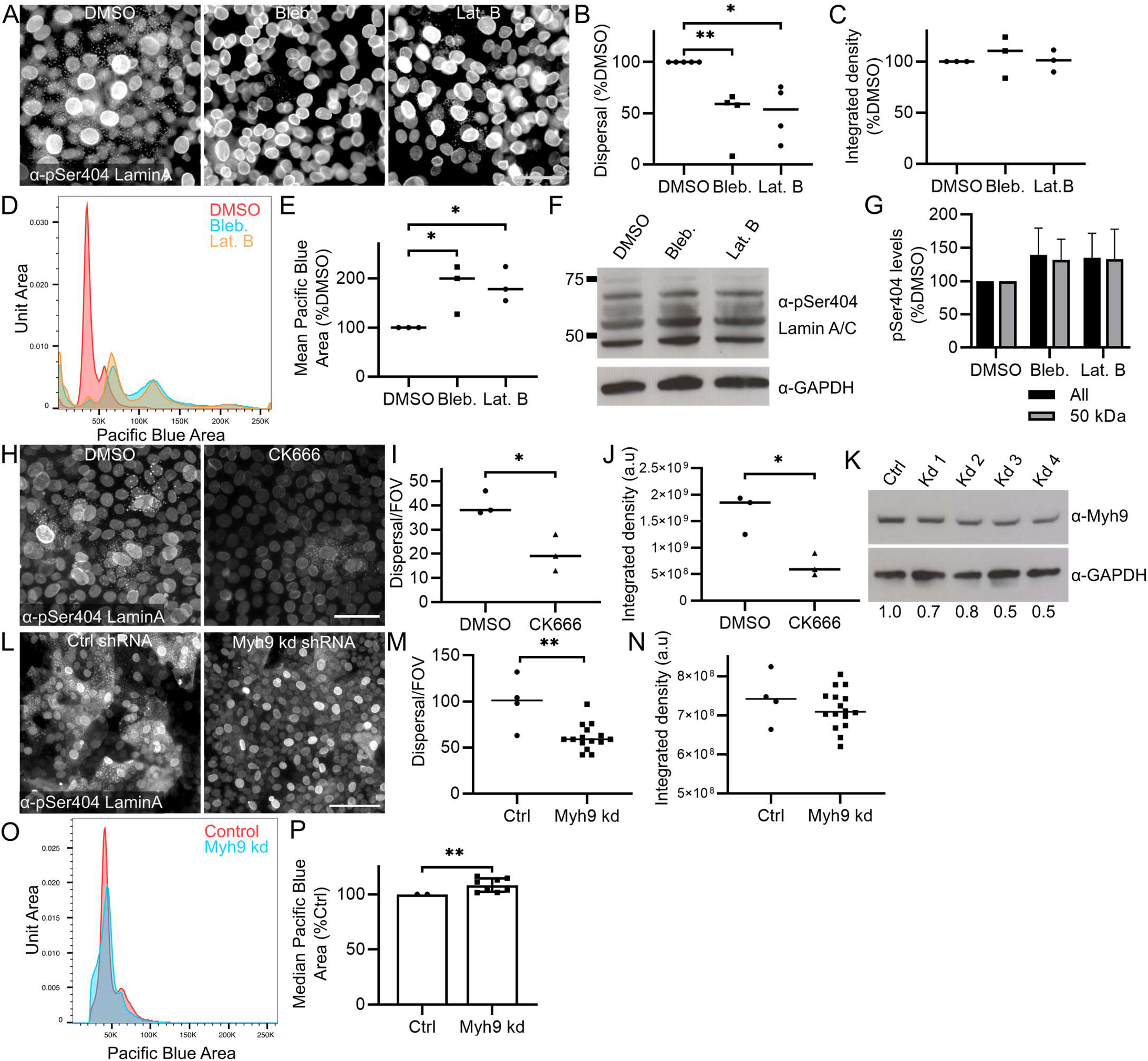
Disruption of cytoskeletal Akt1 target protein function affects nuclear lamina dispersal and nuclear size. A – DMSO, blebbistatin and latrunculin B treated REKs stained for pSer404 Lamin A/C. Epifluorescence images, scale bar = 50 μm. B – Number of cells with pSer404 Lamin A/C dispersal in DMSO, blebbistatin and latrunculin B treated REKs. % of DMSO control, >4 independent experiments, > 3 FOV per experiment, one-way ANOVA * p ≤ 0.05, ** p ≤ 0.01. C – pSer404 Lamin A/C intensity in DMSO, blebbistatin and latrunculin B treated REKs. % of DMSO control, > 3 independent experiments, > 3 FOV per experiment, one-way ANOVA all comparisons non-significant. D – Pacific Blue Area of DMSO, blebbistatin and latrunculin B treated REKs. E - Mean Pacific Blue Area of DMSO, blebbistatin and latrunculin B treated REKs. % of DMSO control, 3 independent experiments, one-way ANOVA, ** p ≤ 0.01, *** p ≤ 0.001. F – pSer404 Lamin A immunoblots of DMSO, blebbistatin and latrunculin B treated REKs. G – pSer404 Lamin A level of all fragments and the ~50 kDa fragment. % of DMSO, 2 independent experiments, all comparisons ns. H – DMSO or CK666 treated REKs stained for pSer404 Lamin A/C. Epifluorescence images, scale bar = 50 μm. I – Number of cells with pSer404 Lamin A/C dispersal in DMSO and CK666 treated REKs. 3 FOV, unpaired t-test * p ≤ 0.05. J – pSer404 Lamin A/C staining intensity in DMSO and CK666 treated REKs. 3 FOV, unpaired t-test * p ≤ 0.05. K - Myh9 expression in Ctrl and Myh9 shRNA knockdown post-confluent REKs. Numbers are relative to GAPDH expression and normalised to Ctrl. L - Ctrl and Myh9 shRNA knockdown post-confluent REKs stained for pSer404 Lamin A/C. Epifluorescence images, scale bar = 50 μm. M – Number of cells with pSer404 Lamin A/C dispersal in Ctrl and Myh9 shRNA knockdown REKs. 4 independent shRNA knockdown cell lines, >3 FOV per line, Mann-Whitney test, ** p ≤ 0.01. Representative of two independent experiments. N - pSer404 Lamin A/C staining intensity in control and Myh9 shRNA knockdown cells. 4 independent shRNA knockdown cell lines, >3 FOV per line, Mann-Whitney test non-significant. Representative of two independent experiments. O – Pacific Blue area of Ctrl or Myh9 shRNA knockdown REKs. P – Median Pacific Blue area in Ctrl and Myh9 shRNA knockdown REKs. 2 independent experiments, 4 independent shRNA knockdown cell lines, Wilcoxon test, ** p ≤ 0.01.

Myh9 (myosin IIa) shRNA knockdown (Figure 5K) also decreased phosphorylated Lamin A/C dispersal, (Figure 5L-N) and increased nuclear size, (Figure 5O-P). This suggested that the activity of myosin II and the presence of the actin cytoskeleton is required for the formation and/or distribution of cytoplasmic Lamin A/C structures after Akt1-dependent phosphorylation of Lamin A/C.

### Fluorescently tagged nuclear proteins illustrate nuclear shrinkage in REKs

To understand how Lamin A/C dispersal occurs in suprabasal differentiating REKs we expressed fluorescently tagged nuclear proteins and imaged them at least three days post-confluency to follow changes in nuclear morphology in real time in differentiating REKs. The majority of EGFP-Lamin A positive REKs did not show any alterations in nuclear size/maximum cross sectional area over a period of 11 hours, (Movie 1). However, 10% of the nuclei rapidly decreased in size, within a period of 40 min, (Movie 2 and Figure 6A and B). There is no further decrease in size after this, (Figure 6C), and this is a decrease in nuclear volume not just due to rotation of the nucleus, (Figure 6D). In post-confluent REK monolayer cultures, DAPI and Lamin A/C staining intensity decreased in suprabasal shrunken nuclei compared to basal ‘non-shrunken nuclei’, conversely, apparent Lamin B1 staining intensity increased in shrunken nuclei, (Figure 5E and F).

**Figure 6.**
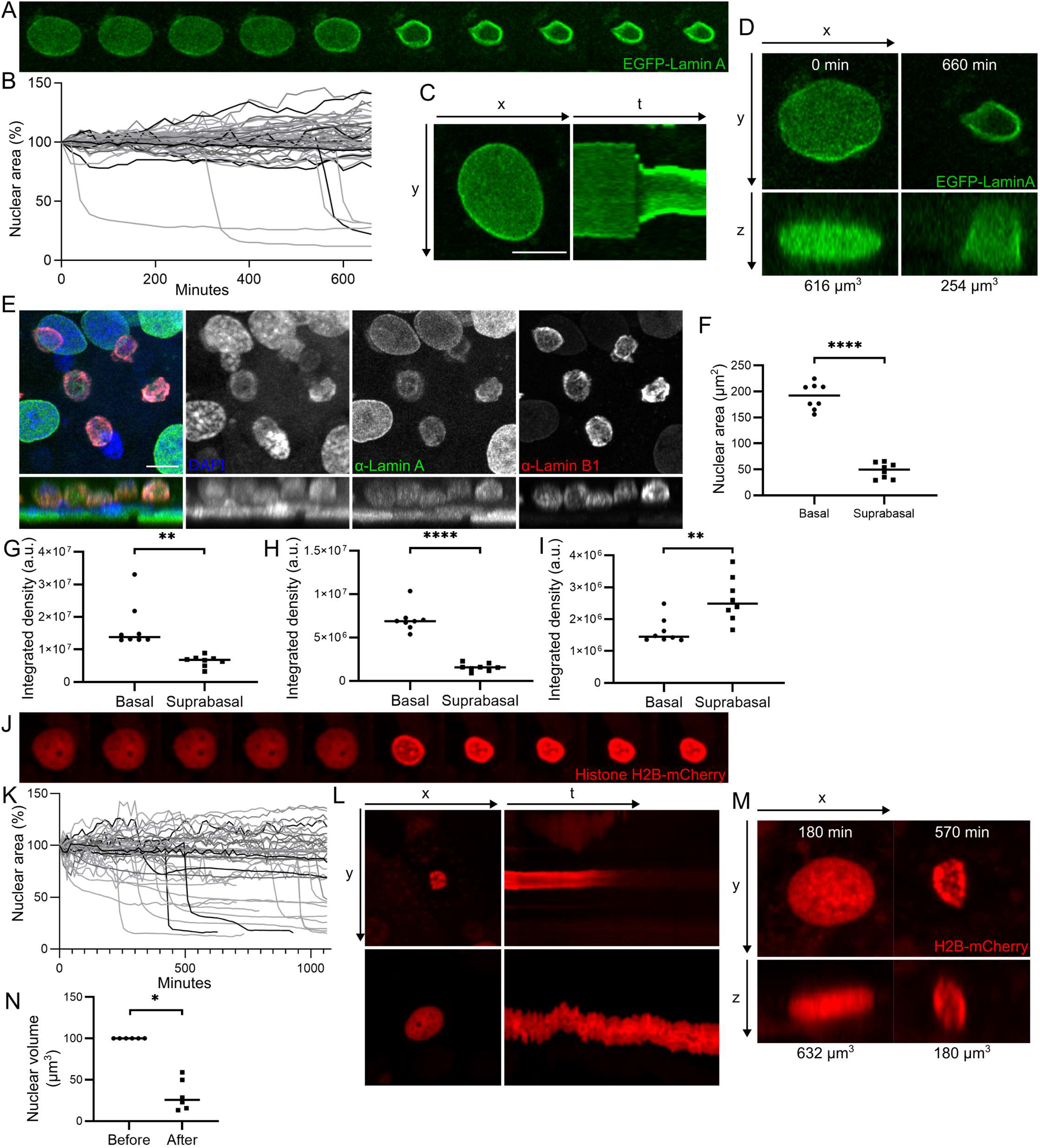
Nuclear dynamics in differentiating REKs. A - Images every 20 min of EGFP-Lamin A positive post-confluent REKs. B – Cross-sectional area EGFP-Lamin A positive nuclei over time. C – Kymographs (yt projection) of a EGFP-Lamin A positive nucleus over time. D – Xz projections of a EGFP-Lamin A positive nucleus. Labelled with nuclear volume. E – Xy and xz projections of confocal z-sections of post-confluent REKs stained for Lamin A and Lamin B1. Scale bar = 10 μm. F ‒Cross-sectional area of basal and suprabasal nuclei in post-confluent REKs. Two independent experiments, Welch’s t-test, **** p ≤ 0.0001. G-I – Intensity of DAPI (G), Lamin A (H) and Lamin B1 (I) staining in basal and suprabasal nuclei in post-confluent REK cultures. Welch’s t test, ** p ≤ 0.01, **** p ≤ 0.0001. Representative of two independent experiments. J - Images every 15 min of Histone H2B-mCherry positive post-confluent REKs. K - Cross-sectional area over time in 45 Histone H2B-mCherry positive nuclei greater than 80 μm^2^. L – Kymograph (yt projection) of Histone H2B-mCherry over time; shrunken nucleus (top panels) and a nucleus that does not shrink (bottom panels). M - Xz projections of Histone H2B-mCherry positive nuclei labelled with nuclear volume. N – Histone H2B-mCherry positive nuclear volume before and after shrinkage. Wilcoxon test, * p ≤ 0.05.

H2B-mCherry expressing REKs also underwent nuclear shrinkage in post-confluent culture. Approximately one quarter of labelled nuclei rapidly decreased in surface area, (Figure 6J-K and Supplementary Figure 3A-C), equating to a decrease in nuclear volume of on average 68.5%, (Figure 6M-N). The intensity of the mCherry signal decreased over time in shrunken nuclei, compared to ‘non-shrunken’ nuclei, until the signal was lost in 41.6% of shrunken nuclei, (Figure 6L and Supplementary Figure 3C). REKs co-expressing Histone H2B-mCherry and EGFP-Lamin A, underwent nuclear shrinkage three days post-confluency with no decrease in mCherry or EGFP signal over 10 hours in shrunken nuclei, (Supplementary Figure 3D-F). Taken together these data indicated that nuclear shrinkage occurs before nuclear content degradation, and maintained expression of Lamin A/C prevented nuclear degradation after nuclear shrinkage.

### Phosphorylated Lamin A/C is cleaved prior to cytoplasmic dispersal

Live imaging of REKs expressing EGFP-Lamin A did not demonstrate dispersal of EGFP-Lamin A to cytoplasmic structures and EGFP-Lamin A did not co-localise with pSer404 Lamin A staining, (Figure 7A). Lamin A/C has been previously predicted to contain two cysteine protease cleavage sites: at Asp230 and Asp446, with cleavage during apoptosis at Asp230 (Rao, Perez, and White 1996; Broers et al. 2002). Single fragments of ~55 kDa in post-confluent EGFP-Lamin A expressing REKs (N-terminal EGFP tag) and ~80 kDa in Lamin A-mEmerald expressing REKs (C-terminal Emerald tag) suggested cleavage of Lamin A at Asp230, (Figure 7B and Supplementary Figure 4A-C). An antibody targeted to the Lamin A/C N-terminus did not stain cytoplasmic pSer404 Lamin A structures, (Figure 7C), indicating that the C-terminal Lamin A/C degradation product, containing the Ser404 phosphorylation site, is targeted to cytoplasmic structures, but not full-length Lamin A/C.

**Figure 7.**
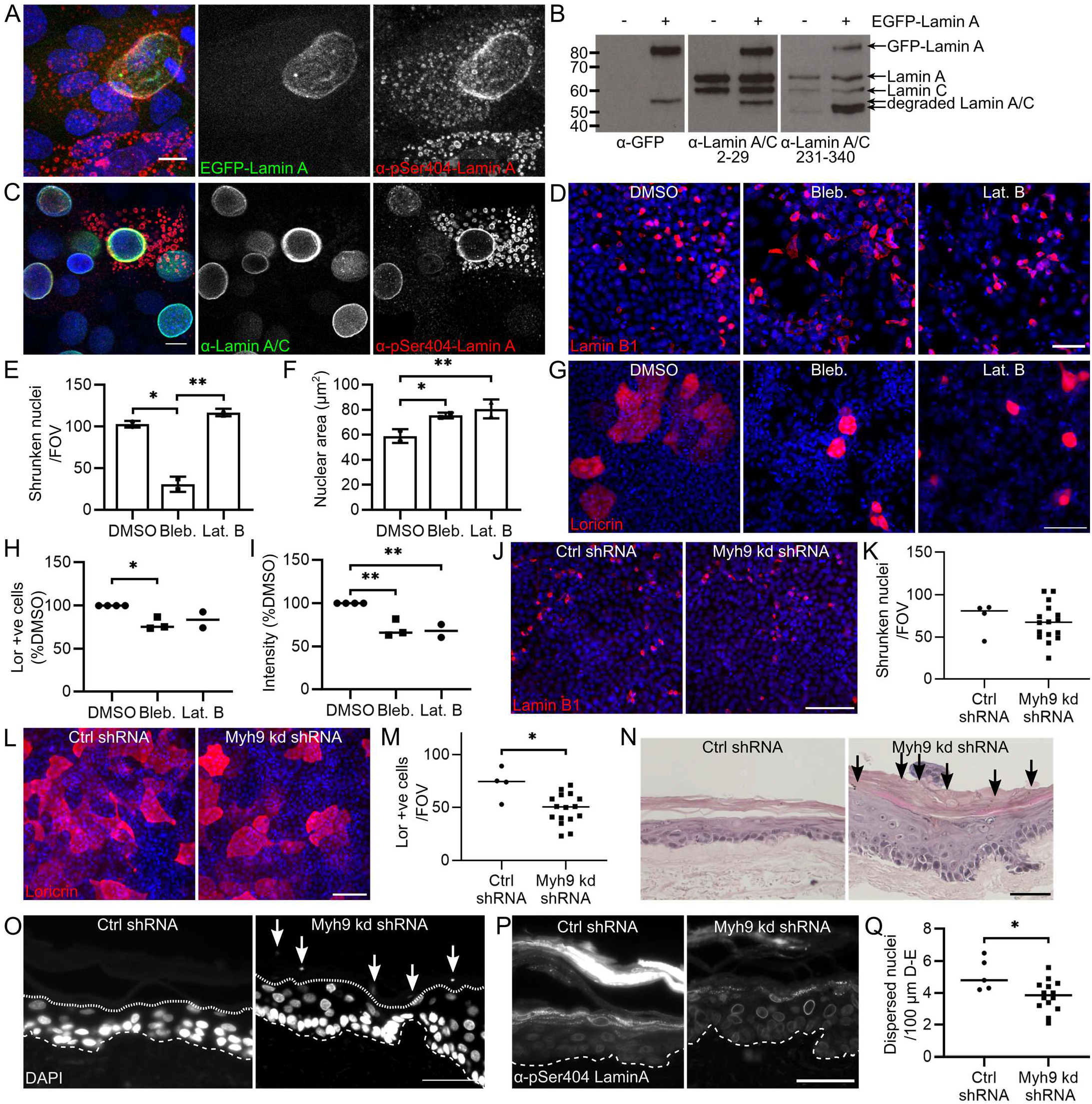
Lamin A cleavage in nuclear degradation and cytoskeletal protein inhibitors prevent nuclear shrinkage. A – EGFP-Lamin A post-confluent REKs stained for pSer404 Lamin A. Maximum projection of confocal z-sections, scale bar = 10 μm. B – Immunoblotting untransfected and EGFP-Lamin A transfected post-confluent REK lysates with antibodies directed to GFP and two Lamin A/C regions. C - Post-confluent REKs stained for pSer404 Lamin A and the N-terminus of Lamin A/C. Single confocal plane, scale bar = 10 μm. D – DMSO, blebbistatin and latrunculin B treated REKs stained for Lamin B1. Scale bar = 50 μm. E-F – Number per FOV (E) and area (F) of shrunken Lamin B1 expressing nuclei in DMSO, blebbistatin and latrunculin B treated REKs. 2 experiments, 3 FOV/experiment, two-way ANOVA, * p ≤ 0.05, ** p ≤ 0.01. G – DMSO, blebbistatin and latrunculin B treated REKs stained for loricrin. Scale bar = 100 μm. H – Number of loricrin positive cells per FOV in DMSO, blebbistatin and latrunculin B treated REKs. >2 experiments, 3 FOV/experiment, one-way ANOVA, * p ≤ 0.05. I – Loricrin staining intensity in DMSO, blebbistatin and latrunculin B treated REKs. % of DMSO control, >2 experiments, 3 FOV/experiment, one-way ANOVA, * p ≤ 0.05. J – Ctrl and Myh9 shRNA knockdown post-confluent REKs stained for Lamin B1. Scale bar = 100 μm. K- Number of shrunken Lamin B1 expressing nuclei per FOV in Ctrl and Myh9 shRNA knockdown post-confluent REKs. 4 independent shRNA knockdown cell lines, >3 FOV per line, unpaired t-test, non-significant. L – Ctrl and Myh9 shRNA knockdown post-confluent REKs stained for loricrin. Scale bar = 100 μm. M - Number of loricrin positive cells per FOV in Ctrl and Myh9 shRNA knockdown post-confluent REKs. 4 independent shRNA knockdown cell lines, >3 FOV per line, Mann-Whitney test, * p ≤ 0.05. N – Ctrl or Myh9 shRNA knockdown REK organotypics stained with hematoxylin and eosin (H&E). Arrows indicate retained nuclei, scale bar = 50 μm. O - Ctrl or Myh9 shRNA knockdown REK organotypics stained with DAPI. Dashed line marks dermal-epidermal junction, dotted line indicates junction of granular and cornified layers and arrows indicate retained nuclei, scale bar = 50 μm. P - Ctrl or Myh9 shRNA knockdown REK organotypics stained for pSer404 Lamin A. Dashed line marks dermal-epidermal junction, scale bar = 50 μm. Q – Number of nuclei with phosphorylated Lamin A/C dispersal per 100 μm of the dermal-epidermal (D-E) junction in Ctrl or Myh9 shRNA knockdown organotypics. 2 independent shRNA knockdown cell lines, 2 organotypics per cell line, >3 FOV per line, Mann-Whitney test, * p ≤ 0.05.

Nuclei expressing a C-terminally tagged Lamin A/C, Lamin A-mEmerald, underwent nuclear shrinkage with the same kinetics as EGFP-Lamin A expressing REKs, (Supplementary Figure 4D-G), but Lamin A-mEmerald also did not disperse, (Supplementary Figure 4H). Lamin A/C contains a C-terminal CAAX box required for farnesylation before prelamin A/C cleavage which is necessary for functional integration of Lamin A/C into the nuclear lamina (Weber, Plessmann, and Traub 1989; Beck, Hosick, and Sinensky 1990; Sinensky et al. 1994). Prevention of CAAX box modification and/or cleavage by presence of the C-terminal mEmerald tag may prevent correct Lamin A-mEmerald integration into the nuclear lamina and consistent with this, the nuclei expressing Lamin A-mEmerald had a disrupted nuclear lamina compared to EGFP-Lamin A expressing nuclei, (Supplementary Figure 4I-J).

### The actomyosin cytoskeleton is required for nuclear shrinkage and epidermal differentiation

We tested the dependence of nuclear shrinkage on the activity of Akt1-associated cytoskeletal proteins in phosphorylated Lamin A/C dispersal. Blebbistatin treatment reduced the number of shrunken Lamin B1 stained nuclei and increased the size of nuclei with high expression of Lamin B1, (Figure 7D-F), indicating a requirement for myosin II activity for both decreased localisation of Lamin A/C at the nucleus and nuclear shrinkage. Latrunculin B treatment also increased the size of nuclei that had high nuclear expression of Lamin B1, (Figure 7D and F), but latrunculin B and CK666 treatment did not affect the overall number of cells in the culture with elevated Lamin B1 expression, (Figure 7E and Supplementary Figure 5A-C). Blebbistatin and latrunculin B treatments caused a reduction in the expression of the epidermal terminal differentiation marker loricrin (Figure 7G-I).

Myh9 shRNA knockdown did not significantly affect nuclear shrinkage, (Figure 7J-K), but did slightly decrease loricrin expression, (Figure 7L-M). Defects in phosphorylated Lamin A/C dispersal and differentiation without defects in nuclear shrinkage may indicate multiple mechanisms that affect nuclear shrinkage beyond Lamin A/C dispersal. Myh9 knockdown organotypic cultures had decreased Myh9 expression, (Supplementary Figure 5D), and were hyperkeratotic and displayed small parakeratotic nuclear material, (Figure 7N-O, Supplementary Figure 5E-F). Fewer cells underwent Lamin A/C dispersal, (Figure 7P-Q), but there was no detectable change in loricrin expression, (Supplementary Figure 5G).

### Lamin A/C dispersal affects granular layer gene expression

To analyse gene expression changes in keratinocytes undergoing differentiation and Lamin A/C dispersal we performed single cell RNA sequencing (RNAseq) on a population of REKs enriched in differentiating keratinocytes. Post-confluent REK cultures were separated by fluorescence-activated flow cytometry into two populations: ‘small’ cells (basal proliferating keratinocytes, with low forward (FSC) and SSC) and ‘large’ cells – (suprabasal differentiating keratinocytes with high FSC and SSC), (Supplementary Figure 5H-I). Mixing these populations at a ratio of 3:1, ‘large’ to ‘small’, yielded a cell suspension with an increased number of differentiating keratinocytes. K-means clustering on single cell RNAseq results delineated two clusters of cells, (Figure 8A), one with higher expression of basal keratins 5 and 14 (K5, K14) and the other with higher expression of suprabasal keratinocyte keratins 1 and 10 (K1, K10), (Figure 8B). These clusters had markedly differential RNA expression, (Figure 8A), and interestingly, the suprabasal cluster had greatly reduced UMI (unique molecular identifier) counts compared to the less differentiated cluster, Figure 8C). Indicating a large reduction in RNA transcripts in suprabasal differentiating keratinocytes. Several transcripts delineated the more differentiated (putative lamin dispersed) population including genes known to be up-regulated during keratinocyte end-stage terminal differentiation including Sprr1a and Krtdap (Kartasova et al. 1988; Tsuchida et al. 2004), (Figure 8D). A number of transcripts, such as those for Rps12 and Tpt1, were enriched in the ‘not dispersed’ population but excluded from the ‘lamin dispersed’ population, (Figure 8D). This mirrored their expression in human epidermis in the Human Protein Atlas (Uhlén et al. 2015), (Figure 8E), indicating their utility as markers of different keratinocyte compartments.

**Figure 8.**
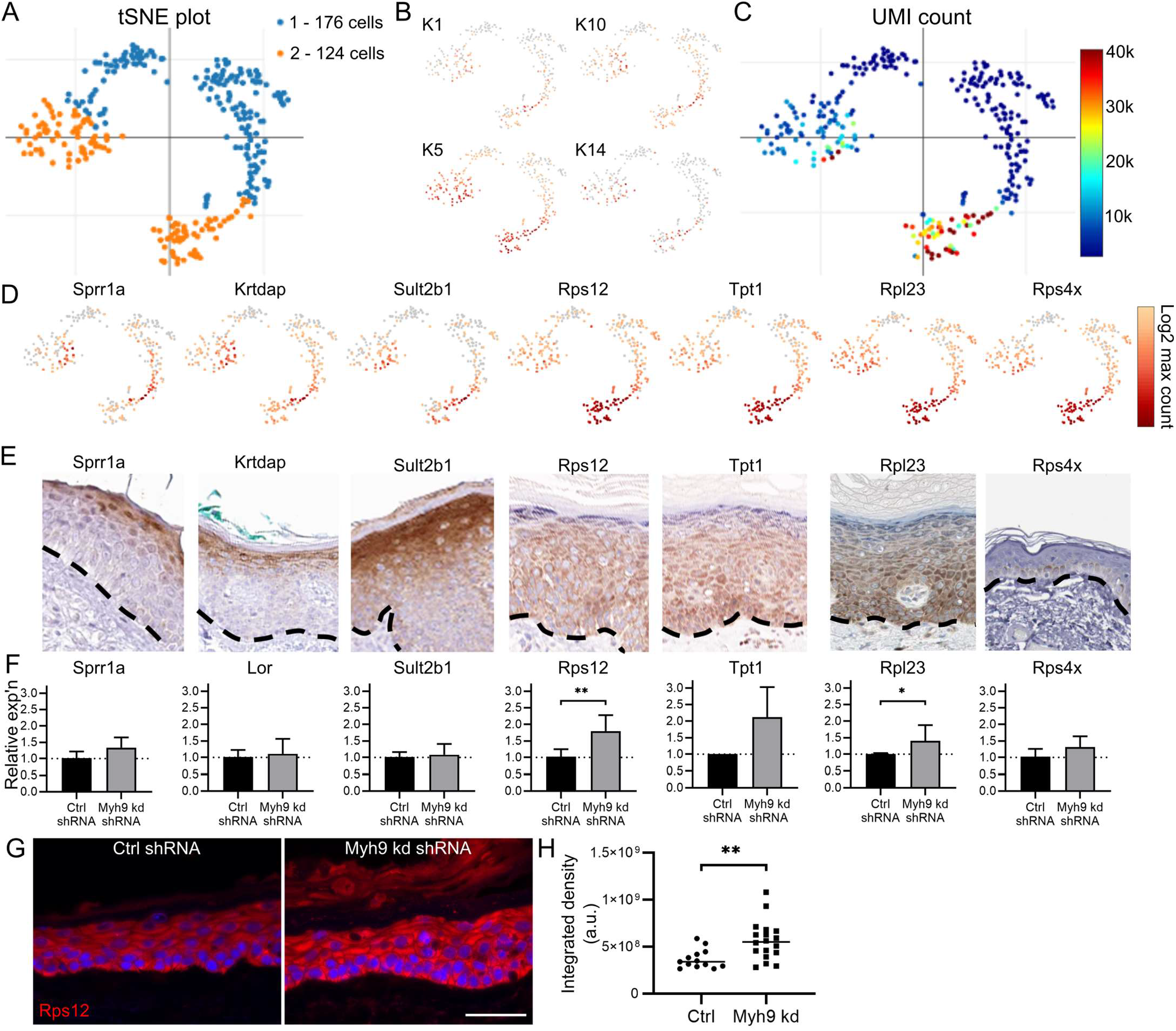
Single cell gene expression in keratinocyte differentiation. A – K-means (K=2) clustering analysis of RNAseq of a differentiation-enriched REK population. B - Expression of basal keratins (K5 and K14) and suprabasal keratins (K1 and K10) in a differentiation-enriched REK population. C – UMI counts (total number of RNA transcripts) of a differentiation-enriched REK population. D – Expression of Sprr1, Krtdap, Sult2b1, Rps12 and Tpt1 in a differentiation-enriched REK population. E – Human epidermal expression of Sprr1, Krtdap, Sult2b1, Rps12 and Tpt1, dermal-epidermal junction indicated by dashed line. Images from the Human Protein Atlas. F – Relative gene expression of Sprr1a, loricrin (Lor), Sult2b1, Tpt1 and Rps12 in Ctrl or Myh9 shRNA knockdown post-confluent REKs. 2 independent experiments, 4 independent shRNA knockdown cell lines, Welch’s t-test, * p ≤ 0.05, ** p ≤ 0.01. G - Ctrl or Myh9 shRNA knockdown REK organotypics stained for Rps12. Scale bar = 50 μm. H – Intensity of Rps12 staining in Ctrl or Myh9 shRNA knockdown REK organotypics. 4 independent shRNA knockdown cell lines, Welch’s t-test, ** p ≤ 0.01.

Myh9 shRNA knockdown increased the expression of genes normally down-regulated in granular layer cells in post-confluent REK cultures, including Tpt1 and Rps12, without up-regulation of differentiation associated genes Sprr1a, Loricrin and Sult2b1, (Figure 8F). Up-regulation of Rps12 at the protein level was also detected in Myh9 shRNA knockdown organotypic cultures, (Figure 8G-H), without up-regulation of loricrin, (Supplementary Figure 5G). This suggests Lamin A/C removal from the nuclear lamina alters gene expression. Blebbistatin treatment additionally increased differentiation marker expression, (Supplementary Figure 5J), consistent with an additional effect caused by its inhibition of other myosin II complexes.

## Discussion

We present data suggesting that nuclear degradation in mammalian epidermal keratinocytes requires Akt1-dependent phosphorylation and cleavage of Lamin A/C before actin cytoskeleton and myosin II mediated cytoplasmic dispersal of pSer404 Lamin A/C. Lamina removal from the nucleus precedes nuclear shrinkage and then nuclear content degradation. Akt1 and its interactors act as part of a ‘degradosome’ complex required for two novel nuclear removal intermediate processes during keratinocyte nuclear destruction: Lamin A/C dispersal and rapid nuclear volume reduction.

Lamins A/C and B1/2 are important components of the nuclear lamina, where they regulate nuclear morphology (Dittmer et al. 2011; Lammerding et al. 2006; Newport, Wilson, and Dunphy 1990; Liu et al. 2000) and control chromatin organisation with the lamin B receptor (Solovei et al. 2013). They interact with DNA through lamina-associated domains (LADs), which are enriched in transcriptionally repressed areas of the genome (Guelen et al. 2008). Various models of lamin association with LADs have been proposed, including direct tethering to chromatin domains (Gonzalez-Sandoval et al. 2015) and a ‘meshwork caging model’ where the dense lamin network traps chromatin domains (Amendola and Steensel 2015; Kim, Zheng, and Zheng 2019). Work in Lamin null (Lamin B1, B1 and A triple knockout) mice demonstrated that loss of LADs altered chromatin organisation and affected gene expression in neighbouring ‘non-LAD’ genomic regions (Zheng et al. 2018). Interestingly, murine skin develops normally without Lamin B1 or B2 expression (Yang et al. 2011) and A and B-type lamins have been suggested to have different functions in chromatin organisation and gene expression regulation (Shevelyov et al. 2009; Solovei et al. 2009; Shevelyov and Ulianov 2019). We demonstrated that Lamin A/C is specifically dispersed to cytoplasmic structures, which do not contain DNA, histone proteins, NLS-targeted protein or Lamin B1. The specific removal of Lamin A/C from the nuclear periphery may specifically alter keratinocyte gene expression to regulate subsequent stages in keratinocyte differentiation.

Our data suggests cleavage of Lamin A/C occurs prior to dispersal from the nucleus, and only the C-terminal fragment of Lamin A/C disperses to the cytoplasm. C-terminally tagged Lamin A-mEmerald was not dispersed, however, the C-terminal tag is likely to affect correct Lamin A/C processing at the C-terminal CAAX box (Sinensky et al. 1994). Progerin, a C-terminal mutant form of Lamin A/C (Eriksson et al. 2003), retains modifications at the CAAX box and is not correctly cleaved, altering nuclear lamina structure and nuclear morphology (Dahl et al. 2006; Lu and Djabali 2018). Therefore, the C-terminal mEmerald tag may prevent correct integration into the nuclear lamina; affect nuclear morphology, Lamin A/C phosphorylation and/or cleavage and dispersal.

We identified myosin IIA activity and the actin cytoskeleton as important for epidermal differentiation and Lamin A/C dispersal. Myosin II activity is required for nuclear repositioning in cell migration (Gomes, Jani, and Gundersen 2005), constriction of the cytokinetic actomyosin contractile ring (Powell 2005; De Lozanne and Spudich 1987; Mabuchi 1977) and contraction of an actin network around the nucleus at nuclear envelope breakdown (Booth et al. 2019). Arp2/3, required for branched actin filament nucleation, has previously been identified as required for epidermal differentiation (Lechler 2014; Zhou et al. 2013), regulation of nuclear actin (Oma and Harata 2011) and correct formation of the actomyosin contractile ring in cell division (Chan et al. 2019). Myosin IIA and Arp2/3 have also been identified as responsible for force generation in the secretion of vesicles in alveolar type II (ATII) cells (Miklavc et al. 2012) and myosin IIA and actin are required for exocytosis in murine exocrine cells (Ebrahim et al. 2019). Disrupting myosin II and Arp2/3 activity affected lamin dispersal, however, whether their activity is important for force generation for formation of cytoplasmic pSer404 Lamin A/C structures or for trafficking of these structures will be important to determine.

Our results suggest another nuclear degradation intermediate of a ‘shrunken nucleus’, which may be the major morphological alteration in nuclear removal. Dispersion of cleaved Lamin A/C without dispersal of nuclear contents prior to nuclear shrinkage suggests that degradation and removal of nuclear lamina proteins may be an initial step in the initiation of nuclear removal. REKs expressing GFP-tagged Lamin A undergo nuclear shrinkage, but do not undergo further degradation. Whereas, REKs expressing Histone H2B-mCherry underwent nuclear shrinkage followed by a gradual decrease in signal intensity, suggesting degradation of nuclear contents. This suggests decreased expression of Lamin A/C at the nuclear lamina is required for nuclear content degradation. Lamin A/C and Lamin B1 form separate but interconnecting networks (Shimi et al. 2008) and perform distinct functions at the nuclear lamina; nuclei deficient for Lamin A/C have decreased nuclear stiffness and increased deformations whereas nuclei deficient for Lamin B1 display normal nuclear mechanics (Lammerding et al. 2006). Protein and lipid depleted nuclear matrices have been shown to reversibly shrink in response to bivalent cation concentration (Wunderlich and Herlan 1977) or nuclease treatment (Berezney and Coffey 1977; Nakayasu and Ueda 1981) and rat nuclei shrink in response to elevated calcium levels (Okada et al. 2010). Degradation of nuclear material occurs after Lamin A/C depletion of the nucleus, as DNase deficient murine epidermis retains DAPI but not Lamin A/C staining in corneocytes (Fischer et al. 2017). Therefore, removal of Lamin A/C may allow subsequent changes in nuclear morphology by altering the stiffness of the lamina, and/or by allowing access for molecules necessary for nuclear content degradation.

Only blebbistatin treatment affected the number of shrunken nuclei in our post-confluent REK cultures. Myh9 (myosin IIA) knockdown, latrunculin B and CK666 treatment did not affect the number of shrunken Lamin B1 expressing nuclei. This suggests nuclear shrinkage may not require Lamin A/C dispersal but that myosin IIB and IIC, which have some partially redundant properties but distinct cellular localisations (Pecci et al. 2018; Sandquist and Means 2008; Wang et al. 2010; Jiang, Kolpak, and Bao 2010; Wylie and Chantler 2008), may be important for nuclear shrinkage.

Single cell RNA sequencing of a REK population enriched for differentiating cells demonstrated a massive decrease in the quantity of RNA expression in differentiating REKs and identified genes that were present or absent from more differentiated REKs. Myh9 knockdown, which decreases Lamin A/C dispersal, up-regulated expression of genes normally down-regulated at the granular layer of keratinocyte differentiation. The organisation of sub-nuclear domains alters in keratinocyte differentiation, with changes in the arrangement of heterochromatin (Gdula et al. 2013) and in progeria, expression of incorrectly modified mutant Lamin A alters chromatin organisation and LAD association with the nuclear lamina (Goldman et al. 2004; Shumaker et al. 2006; McCord et al. 2013). This indicates that Lamin A/C removal from the nucleus may be important for changes in gene expression in keratinocyte differentiation, particularly, for preventing expression of genes normally down-regulated in granular layer cells through alterations to LADs.

Targeted autophagy of the nucleus, nucleophagy, is required for keratinocyte nuclear removal; with localisation of LC3 and autophagosomes to indentations at the nuclear periphery of differentiating keratinocytes (Akinduro et al. 2016). Additionally, impaired clearance of nuclear material by autophagosomes was identified in laminopathies, diseases with mutations in lamina proteins (Park et al. 2009), suggesting that alterations to nuclear lamina structure are key for nuclear destruction. We suggest that dispersal of pSer404 Lamin A/C to cytoplasmic structures and nucleophagic processes at the nuclear periphery may act concurrently to complete nuclear removal.

Identification of the molecular mechanisms that initiate and are involved in nuclear removal would be important for understanding and treating parakeratotic skin diseases. However, this could also be of wider importance in other cell types to prevent cellular processes including proliferation and to activate cell death by programmed nuclear clearance. We hope that further delineation of the processes leading to nuclear removal in healthy epidermis will advance this aim.

## Author contributions

CR and RO conceived and designed the experiments. CR, DW, KO, SO and ALM performed experiments. CR and RO wrote the manuscript.

## Acknowledgements

We would like to thank our colleagues from Queen Mary University of London: Dr Jan Soetaert and Dr Belén Martín-Martín from the Blizard Advanced Light Microscopy Core Facility for help with live imaging, Dr Gary Warnes from the Blizard Institute Flow Cytometry Core Facility for help with flow cytometry and cell sorting, Bart’s and the London Genome Centre for single cell RNA sequencing, Dr Vinni Rajeeve from the CRUK Barts Centre Mass Spectrometry Facility for mass spectrometry and Dr John Connelly for providing blebbistatin and latrunculin B.

## Methods

### Antibodies and materials

The following commercially available antibodies were used: Actin (Sigma-Aldrich, A2066), β-Actin (Millipore, MAB1501R), Akt1 (Cell Signaling Technology, 2967), Arp3 (Santa Cruz Biotechnology, sc-48344), COPS4 (Invitrogen, PA5-57863), FAK1 (Abgent, AP7715a), FLAG (Sigma-Aldrich, F3165), GAPDH Millipore Sigma MAB374, GFP (Santa Cruz Biotechnology, sc-9996), Histone H2B (Santa Cruz Biotechnology, sc-515808), Histone H3.1 (Santa Cruz Biotechnology, sc-517576), HspB1 (Abcam, ab12351), Jup (Santa Cruz Biotechnology, SC-514115), K10 (BioLegend, 905401), Lamin A (Santa Cruz Biotechnology; sc-376248, sc-398927, sc-518013), Lamin B1 (Abcam, ab16048), Loricrin (BioLegend, 905101), Myh9 (GeneTex, GTX113236), Plectin (Santa Cruz Biotechnology, sc-33649), Ran (Santa Cruz Biotechnology, sc-271376) and STAMBP (Biorbyt, ORB341234). An antibody to pSer404 of Lamin A was raised in rabbit to phosphopeptide CGRASp-SHSSQTQGGG by Mimotopes (UK) Ltd, and another was a gift from Sandra Marmiroli (Figure 1C, 2E, 7A and Supplementary Figure 1G). Alexa Fluor 568 Phalloidin was from Thermo Fisher Scientific. EGFP-Lamin A was a gift from Pekka Taimen. Lamin A-mEmerald (mEmerald-LaminA-N-18) and NLS-mCherry (mCherry-Nucleus-7) were gifts from Michael Davidson (Addgene plasmids #54139 and #55110, unpublished). Histone H2B-mCherry was a gift from Robert Benezra (Addgene plasmid #20972, (Nam and Benezra 2009)). S404A, S404D and WT Lamin A constructs were a gift from Sandra Marmiroli. shRNA constructs were obtained from QIAGEN (SureSilencing Akt1 shRNA plasmids) or Origene (HuSH Myh9 shRNA constructs).

### Cell culture

REKs were cultured in DMEM supplemented with 10% FBS, 1% penicillin/streptomycin at 37 °C and 5% CO_2_. For transfection, cells were transfected with Lipofectamine 2000 (Invitrogen) or jetOptimus (Polyplus transfection) according to manufacturers’ guidelines one day after seeding. After transfection with Akt1 or Myh9 shRNA plasmids, knockdown REKs were selected for by addition of G418 (400 μg/ml) or puromycin (1.25 μg/ml), respectively, to the medium for 10 days. Upon confluency REKs were fed every day and cultures fixed 2-4 days post confluency to analyse differentiation.

### Western blotting

Cells were lysed in total lysis buffer (20% β-mercaptoethanol, 5% sodium dodecyl sulphate (SDS), 10 mM Tris pH 7) and boiled for 10 min at 95°C. Protein lysates were separated on 4-20% SDS-polyacrylamide gels (Bio-Rad) and transferred to nitrocellulose membrane before blocking and with 5% milk (Marvel) or bovine serum albumin (Sigma) w/v in PBS-T (0.1% Tween-20 in PBS). The blocked membrane was probed with primary and HRP-linked secondary antibodies (Dako) diluted in the blocking buffer. Luminol (Santa Cruz Biotechnology) was used to detect protein bands and results were analysed with Fiji (Schindelin et al. 2012).

### Co-immunoprecipitation

Cells were lysed in RIPA buffer (50 mM Tris-HCl pH 7.4, 150 mM NaCl, 1% Triton X-100, 0.1% sodium deoxycholate, 0.5% SDS) and immunoprecipitation carried out using the Dynabeads Protein G Immunoprecipitation Kit (Thermo Fisher Scientific) according to the manufacturer’s instructions. Eluted proteins and controls were analysed by western blotting or Coomassie Blue staining (Fisher).

### Immunocytochemistry

REK monolayer cultures were fixed in 4% PFA 0.2% triton X-100 or −20 °C methanol. Blocking was performed with 0.4% fish skin gelatin in PBS with 0.2% Triton X-100 and primary and Alexa fluor-conjugated secondary antibodies were diluted in blocking solution. Samples were counterstained with DAPI, mounted with Prolong Gold (Life Technologies) and imaged on Leica DM5000B epifluorescence and Zeiss LSM880 Airyscan confocal microscopes. Analysis was performed in Fiji; pSer404 Lamin A dispersing cells were counted using the ImageJ Cell Counter plugin, Lamin B1 shrunken nuclei were thresholded and then counted using the Analyze Particles plugin.

### Fluorescent immunohistochemistry

Wild-type neonatal murine sections (5 μm) were dewaxed and antigens retrieved in boiling 0.01 M sodium citrate pH 7 for 7 min. 0.4% fish skin gelatin 0.2% Triton X-100 in PBS blocking buffer was also used for primary and secondary antibody dilutions. Samples were mounted with Prolong Gold with DAPI (Life Technologies) and imaged on Leica DM5000B epifluorescence and Zeiss LSM880 Airyscan confocal microscopes.

### Live imaging

REKs transfected with fluorescently tagged proteins were seeded in CELLview™ Cell Culture Dishes and allowed to reach confluency and then cultured for a further 3-4 days before imaging. Confocal images were taken every 15/20 min on a Zeiss LSM880 Airyscan confocal microscope. Images were processed and analysed using Fiji.

### Flow cytometry and cell sorting

Post-confluent REK cultures were dissociated in trypsin, resuspended in PBS with Hoechst 33342 and incubated for 45 min at 37 °C. Flow cytometry was performed on a FACS Canto II (BD Biosciences) and analysed with FlowJo (BD). Cells were sorted, without Hoechst incubation, on a FACSAria IIIu Cell Sorter (BD Biosciences).

### Mass spectrometry

Collodial coomassie dye stained acrylamide gel bands were destained and digested using the method as reported (Shevchenko et al. 2007). The extracted peptides were further cleaned using C18+carbon top tips (Glygen corporation, TT2MC18.96) and eluted with 70% acetonitrile (ACN) with 0.1% formic acid.

Dried peptides were dissolved in 0.1% TFA and analysed by nanoflow ultimate 3000 RSL nano instrument was coupled on-line to a Q Exactive plus mass spectrometer (Thermo Fisher Scientific). Gradient elution was from 3% to 35% buffer B in 120 min at a flow rate 250nL/min with buffer A being used to balance the mobile phase (buffer A was 0.1% formic acid in water and B was 0.1% formic acid in ACN). The mass spectrometer was controlled by Xcalibur software (version 4.0) and operated in the positive mode. The spray voltage was 1.95 kV and the capillary temperature was set to 255 °C. The Q-Exactive plus was operated in data dependent mode with one survey MS scan followed by 15 MS/MS scans. The full scans were acquired in the mass analyser at 375-1500m/z with the resolution of 70 000, and the MS/MS scans were obtained with a resolution of 17 500.

MS raw files were converted into Mascot Generic Format using Mascot Distiller (version 2.5.1) and searched against the SwissProt database restricted to Rat entries using the Mascot search daemon (version 2.5.0) with a FDR of ~1% and restricted to the human entries. Allowed mass windows were 10 ppm and 25 mmu for parent and fragment mass to charge values, respectively. Variable modifications included in searches were oxidation of methionine, pyro-glu (N-term) and phosphorylation of serine, threonine and tyrosine. The mascot result (DAT) files were extracted into excel files for further normalisation and statistical analysis.

### Single cell RNA sequencing

REKs were sorted into ‘small’ – low FSC and SSC and ‘large’ – high FSC, populations and these were mixed at a ratio of 1:3 to generate a population with an enriched number of differentiating cells. For sample quality control, the single cell suspension was assessed for cell number using the Luna FL automated cell counter (Logos biosystems, South Korea). Cells appeared intact and well distributed with a total count of 250 cells/μL. An equivalent volume of 400 cells was loaded to the 10X Chromium™ Single Cell A Chip (PN-1000009) using the Chromium™ 3’ Library & Gel Bead Kit v2 (PN-120267) as described in the manufacturers user guide (10X Genomics, California, USA). GEMs were recovered from the chip and appeared opaque and uniform in colour. 14 cycles of cDNA amplification were performed on the purified GEM-RT product, and cDNA was examined for quality using the Agilent 2200 Tapestation with the High-sensitivity D5000 screentape and reagents (Agilent Technologies, Waldbronn, Germany), and the Qubit^®^ 2.0 Fluorometer and Qubit dsDNA HS Assay Kit (Life Technologies, California, USA). 163ng of cDNA was used to prepare the 10X 3’RNA library and 12 cycles were used for sample index PCR. Final cleaned libraries were quantified using the Qubit^®^ 2.0 Fluorometer and Qubit dsDNA HS Assay Kit and average fragment size checked using the Agilent D1000 screentape and reagents. Final library was run on a 150-cycle v3 MiSeq kit with a 26[8]98 cycle configuration to generate 10 million read pairs in total. Raw sequence data was processed using the 10X Genomics cellranger pipeline(v3.0.2). Briefly, fastq files were generated for the sample, followed by barcode processing and alignment to the Rnor6 genome reference using cellranger count. 178 barcodes were identified as cells. K-means cluster analysis and gene expression analysis was performed in Loupe Cell Browser.

