## Supplementary Figures for "Akt1-associated actomyosin remodelling is required for nuclear lamina dispersal and nuclear shrinkage in epidermal terminal differentiation"

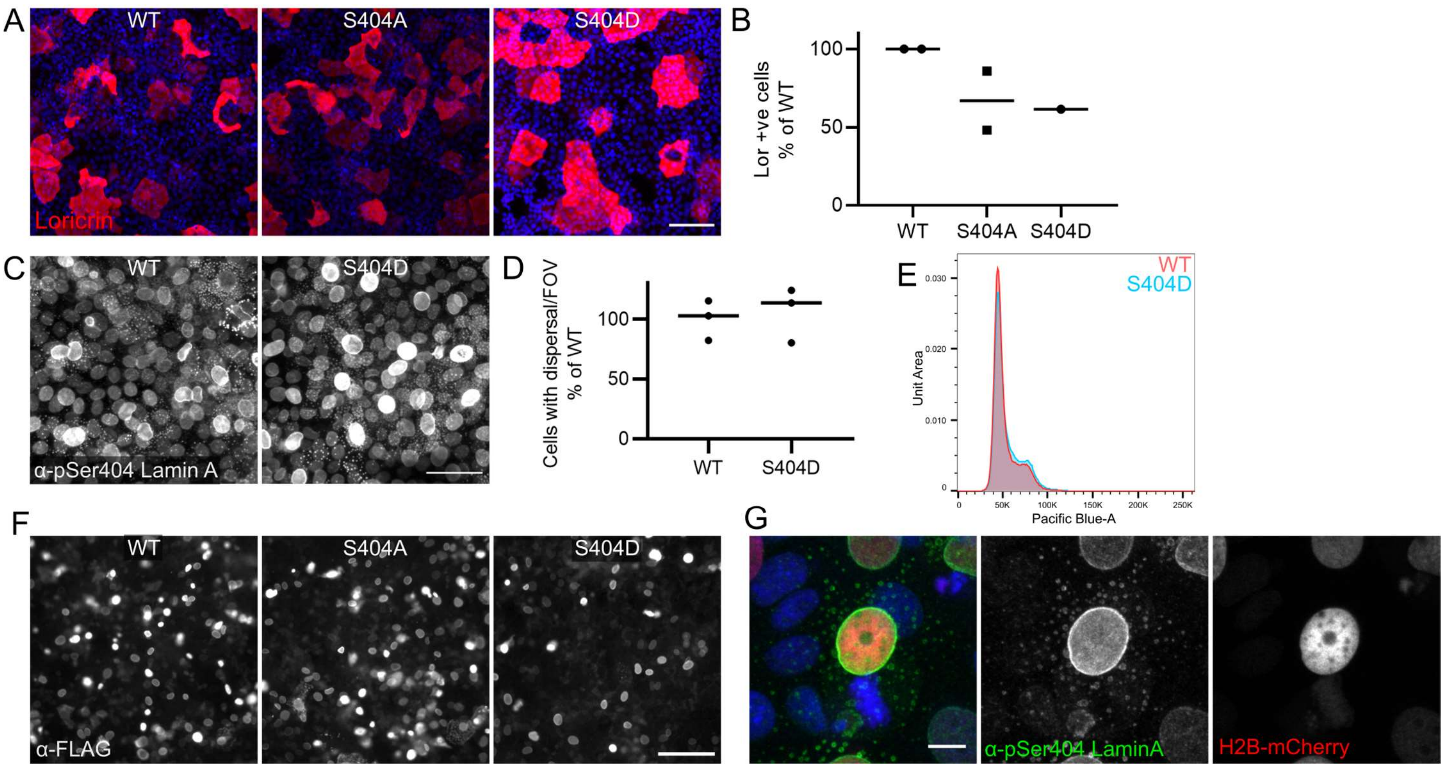


**Supplementary Figure 1 – pSer404 Lamin A dispersal and loricrin expression in REKs.**

A – Post-confluent REK cultures expressing WT, S404A or S404D Lamin A/C stained for loricrin. Scale bar = 100 µm.

B – Number of loricrin expressing cells in REK cultures expressing WT, S404A or S404D Lamin A/C. % of WT, > 3 FOV per experiment, one-way ANOVA, non-significant.

C – Post-confluent REK cultures expressing WT or S404D Lamin A/C stained for pSer404 Lamin A/C. Scale bar = 50 µm.

D – Number of cells with dispersed pSer404 Lamin A/C in post-confluent REK cultures expressing WT or S404D Lamin A/C. % of WT, 3 FOV per construct, unpaired t-test, non-significant.

E - Area of Hoechst 33342 staining of WT or S404D Lamin A/C expressing REKs.

F – Post-confluent REK cultures expressing WT, S404A or S404D Lamin A/C stained for FLAG. Scale bar = 100 µm.

G – Histone H2B-mCherry positive post-confluent REKs stained for pSer404 Lamin A/C. Maximum projection of confocal z-sections, scale bar = 10 µm.

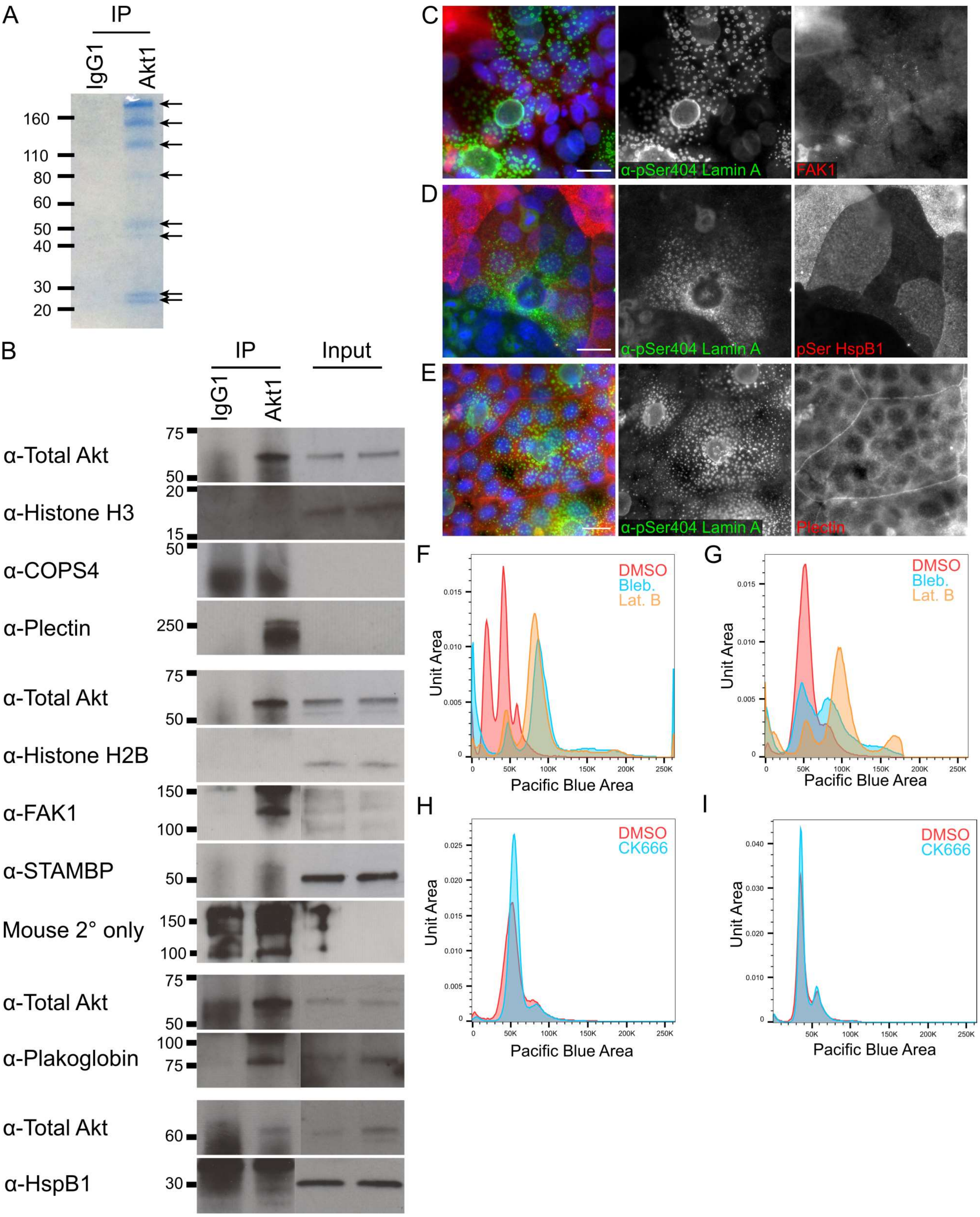


**Supplementary Figure 2 –Akt1 interactor identification and inhibition of cytoskeletal candidates is required for nuclear size.**

A – Coomassie staining of IgG1 and Akt1 immunoprecipitate (IP).

B – IgG1 and Akt1 IPs immunoblotted for Histone H3, COPS4, Plectin, Histone H2B, FAK1, STAMBP, Plakoglobin and HspB1.

C-E - Post-confluent REKs co-stained for pSer404 Lamin A/C and FAK1 (C), pSer86 HspB1 (D) and Plectin (E). Epifluorescence images, scale bars = 20 µm.

F – Area of Hoechst 33342 signal of DMSO, blebbistatin and latrunculin B treated REKs, replicate experiment.

G - Area of Hoechst 33342 signal of DMSO, blebbistatin and latrunculin B treated REKs, replicate experiment.

H - Area of Hoechst 33342 signal of DMSO and CK666 treated REKs, replicate experiment.

I - Area of Hoechst 33342 signal of DMSO and CK666 treated REK cultures, replicate experiment.

A

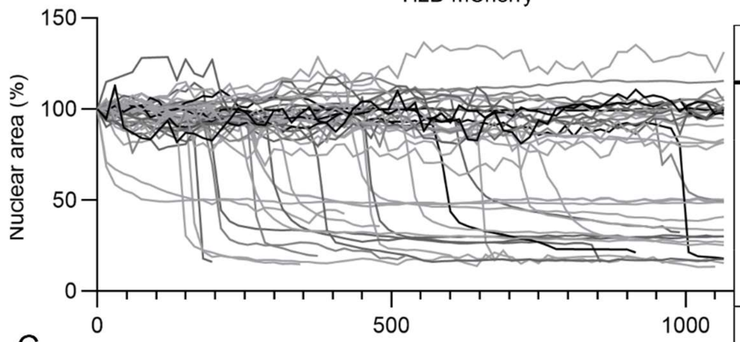


B

| Initial size | No. (%) | Shrink in imaging | No. (%) | Fade in imaging | No. (%) |
|---|---|---|---|---|---|
| Large | 222 (74.2%) | Yes | 42 (18.9%) | Yes | 22 (52.4%) |
| | | | | No | 20 (47.5%) |
| | | No | 180 (81.1%) | Yes | 14 (7.8%) |
| | | | | No | 166 (92.2%) |
| Shrunken | 77 (25.8%) | Yes | 0 | - | - |
| | | No | 77 (100%) | Yes | 32 (41.6%) |
| | | | | No | 45 (58.4%) |

C

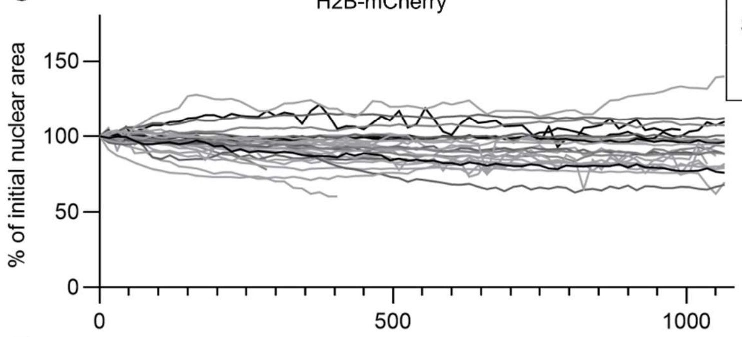


D

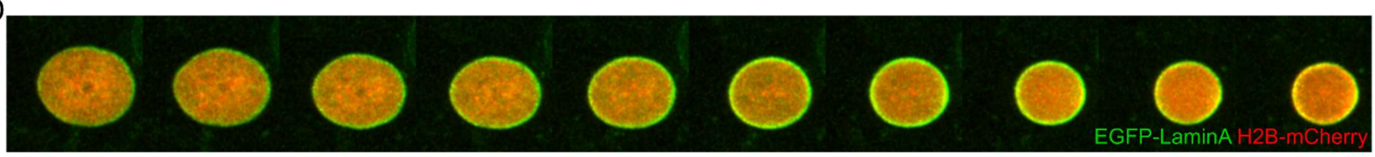


E

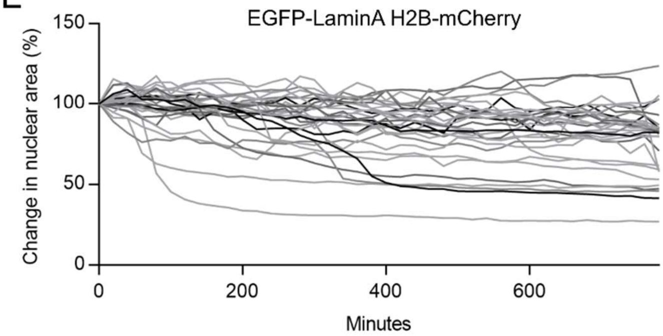


F

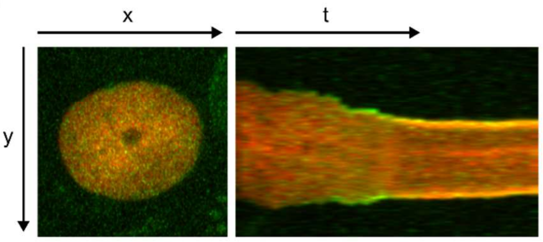


**Supplementary Figure 3 – Live imaging of nuclei positive for Histone H2B-mCherry alone and with EGFP-Lamin A.**

A - Cross-sectional area over time of a further 44 Histone H2B-mCherry positive nuclei greater than 80 μm$^2$.

B – Number of Histone H2B-mCherry positive nuclei that shrink and fade during imaging.

C - Cross-sectional area over time of Histone H2B-mCherry positive nuclei less than 80 μm$^2$.

D - Images every 20 min of a Histone H2B-mCherry and EGFP-Lamin A positive nucleus in post-confluent REKs.

E - Cross-sectional area over time of Histone H2B-mCherry and EGFP-Lamin A positive nuclei.

F – Kymograph (yt) of a Histone H2B-mCherry and EGFP-Lamin A expressing nucleus.

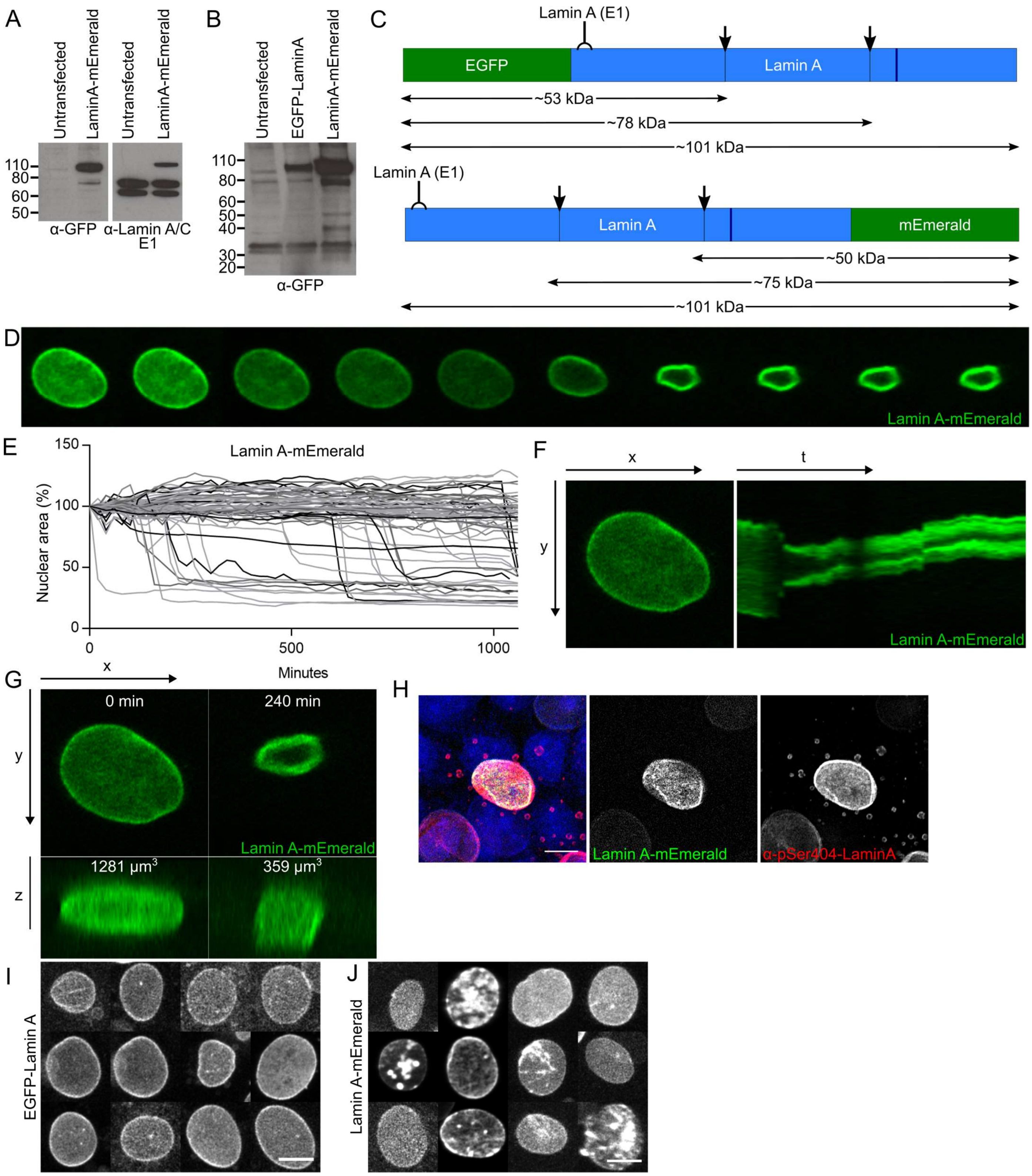


**Supplementary Figure 4 – GFP-tagged Lamin A dynamics in post-confluent REKs.**

A – Untransfected or Lamin A-mEmerald expressing post-confluent REK lysates immunoblotted for GFP or the N-terminus of Lamin A/C (E1).

B – Overexposed blot of untransfected, EGFP-Lamin A or Lamin A-mEmerald expressing post-confluent REK lysates immunoblotted for GFP.

C - Diagram of Lamin A primary protein structure, highlighting predicted cleavage sites (orange lines and arrows), Ser404 (blue), N-terminal antigen for antibody (E1), relative to N-terminal or C-terminal tags.

D - Images every 20 min of a Lamin A-mEmerald positive nucleus.

E - Cross-sectional area over time of Lamin A-mEmerald positive nuclei.

F – Kymograph (yt) of a Lamin A-mEmerald positive nucleus.

G - Xz projections of a Lamin A-mEmerald positive nucleus. Labelled with nuclear volume.

H – Lamin A-mEmerald positive REKs stained for pSer404 Lamin A. Confocal z-section, scale bar = 10 µm.

I – Morphology of EGFP-Lamin A positive nuclei in the first 30 min of imaging, 12 nuclei randomly selected from 12 FOV. Sum of confocal z-sections, scale bar = 10 µm.

J – Morphology of Lamin A-mEmerald positive nuclei in the first 30 min of imaging, 12 nuclei randomly selected from 12 FOV. Sum of confocal z-sections, scale bar = 10 µm.

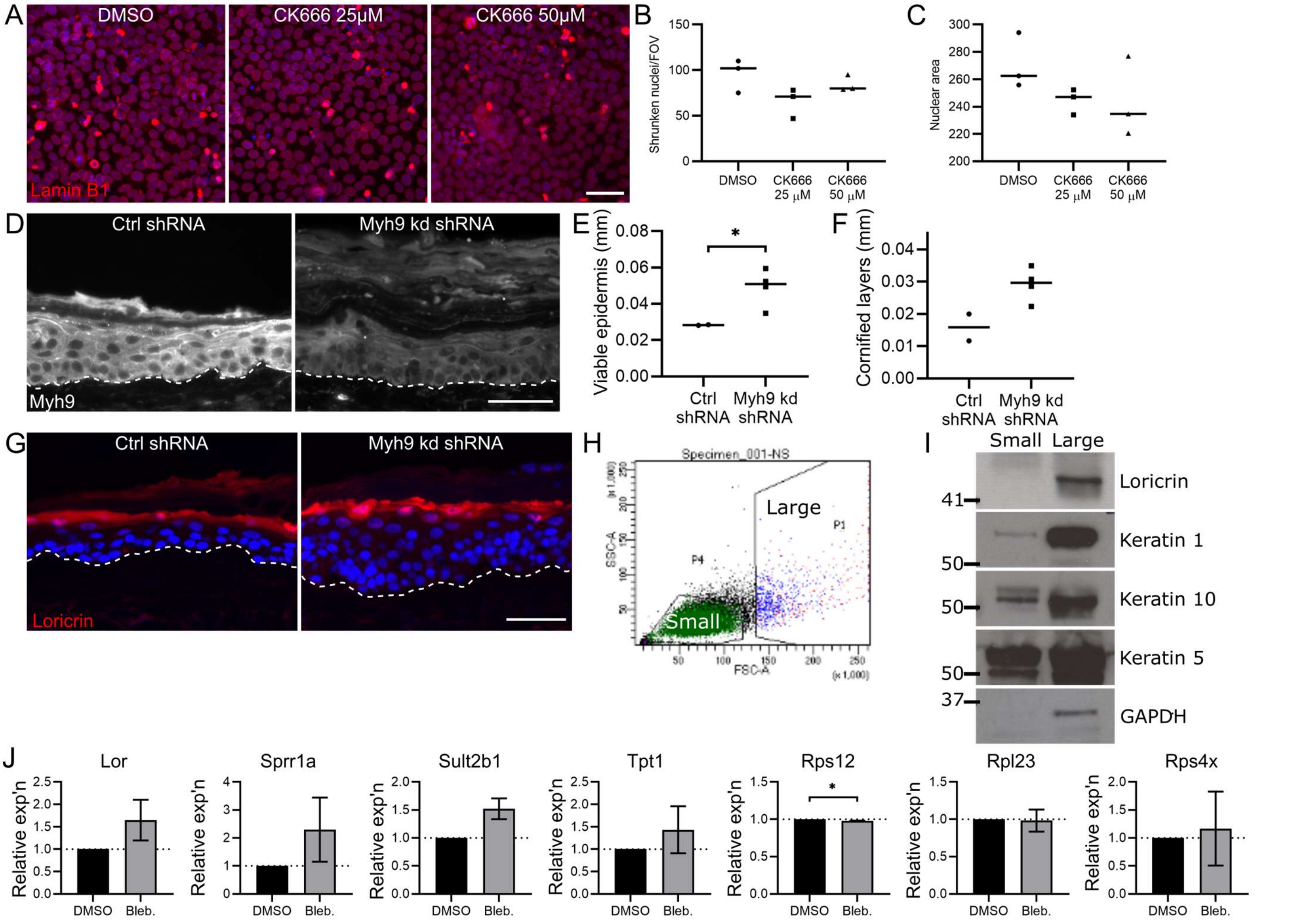


**Supplementary Figure 5 – Arp3 inhibitor CK666 treatments, cell sorting data and relative gene expression in Myh9 knockdown and blebbistatin treated experiments.**

A – DMSO and CK666 treated REKs stained for Lamin B1. Scale bars = 50 µm.

B - Number of shrunken Lamin B1 expressing nuclei per FOV in DMSO and CK666 treatments. Three fields of view, all comparisons non-significant.

C – Nuclear area of shrunken Lamin B1 expressing nuclei in DMSO and CK666 treatments. Three fields of view, all comparisons non-significant.

D - Ctrl and Myh9 shRNA knockdown organotypics stained for Myh9. Scale bar = 50 µm.

E-F – Height of viable epidermis (E) and cornified layers (F) in H&E stained Ctrl and Myh9 shRNA knockdown organotypics. 2 organotypics per line, two independent Myh9 shRNA knockdown lines, >3 measurements per FOV, Welch's t test, * $p \leq 0.05$.

G - Ctrl and Myh9 shRNA knockdown organotypics stained for loricrin. Scale bar = 50 µm.

H – Scatter plot from sorting 'small' basal cells and 'large' differentiating cells from post-confluent REKs using FSC and SSC.~

I – Sorted 'small' and 'large' REK population lysates immunoblotted for loricrin, keratin 1, 10, 5 and GAPDH. ~14% more 'small' cells loaded than 'large' cells.

J - Relative gene expression of loricrin, Sprr1a, Sult2b1, Tpt1, Rps12, Rpl23 and Rps4x in DMSO and blebbistatin treated REKs. 2 independent experiments, one-sample t-test, * $p \leq 0.05$.
